# Hemagglutinin stability determines influenza A virus susceptibility to a broad-spectrum fusion inhibitor Arbidol

**DOI:** 10.1101/2022.01.11.475941

**Authors:** Li Zhenyu, Li Tian, Liu Meisui, Ivanovic Tijana

## Abstract

Understanding mechanisms of resistance to antiviral inhibitors can reveal nuanced features of targeted viral mechanisms and, in turn, lead to improved strategies for inhibitor design. Arbidol is a broad-spectrum antiviral which binds to and prevents the fusion-associated conformational changes in the trimeric influenza hemagglutinin (HA). The rate-limiting step during HA-mediated membrane fusion is the release of the hydrophobic fusion peptides from a conserved pocket on HA. Here, we investigated how destabilizing or stabilizing mutations in or near the fusion peptide affect viral sensitivity to Arbidol. The degree of sensitivity was proportional to the extent of fusion peptide stability on the pre-fusion HA: stabilized mutants were more sensitive, and destabilized ones resistant to Arbidol. Single-virion membrane fusion experiments for representative Wild Type and mutant viruses demonstrated that resistance is a direct consequence of fusion-peptide destabilization not dependent on reduced Arbidol binding to HA at neutral pH. Our results support the model whereby the probability of individual HAs extending to engage the target membrane is determined by the composite of two critical forces: a ‘tug’ on the fusion peptide by the extension of the central coiled-coil on HA, and the key interactions stabilizing fusion peptide in the pre-fusion pocket. Arbidol increases the free-energy penalty for coiled-coil extension, but destabilizing mutations decrease the free-energy cost for fusion peptide release, accounting for the observed resistance. Our findings have broad implications for fusion-inhibitor design, viral mechanisms of resistance, and our basic understanding of HA-mediated membrane fusion.

## Introduction

To deliver their infectious cargo to the host cell, enveloped viruses mediate fusion between the viral and the target cell membranes. A homotrimeric glycoprotein, hemagglutinin (HA), serves as a catalyst for influenza A virus (IAV) membrane fusion. HA is synthesized as an inactive precursor HA0 and becomes activated for fusion by cleavage into the disulfide-linked HA1 and HA2 subunits(1). HA1 forms the globular head domain harboring the receptor binding domain, and serves to stabilize the pre-fusion conformation of HA2. HA2 forms the highly conserved stem region and mediates membrane fusion by undergoing large-scale conformational changes triggered by low pH in late endosomes(2). Proton binding by HA1/2 triggers HA1 head opening, which releases the constraints on HA2 conformational changes (3). The rate-limiting step in HA-mediated membrane fusion is the release of hydrophobic peptides on the HA2 N termini from their buried location near the trimer axis at the base of HA (4).The extension of the central coiled coil on HA2 projects the fusion peptides away from the viral membrane and toward the target membrane where they can insert (5). A single inserted HA cannot overcome the hydration force barrier to membrane fusion (4, 6, 7), but depending on virion size, tens to thousands of HAs interface the target membrane. If several neighboring HAs (fusion cluster) insert their fusion peptides into the target membrane, they cooperate to bring about membrane fusion (4). Mixing of the outer viral and endosomal membrane leaflets in a hemifusion intermediate precedes pore opening and full fusion (8). Previous single-virion membrane fusion experiments defined the kinetics but not the sequence of HA conformational changes leading to hemifusion (4, 8–10). It remains an open question whether HA coiled-coil extension drives fusion-peptide release, or whether fusion-peptide release precedes and enables HA coiled-coil extension.

Inhibition of viral membrane fusion is an effective strategy of inhibiting viral infection. Antibodies and small-molecule inhibitors targeting HA, or weak bases that raise endosomal pH can all be effective at inhibiting viral membrane fusion (11–16). However, resistance to antivirals remains inevitable, and understanding both the mechanism of inhibitor action and resistance phenotypes is essential for improving inhibitor efficacy and devising strategies for extending their effects. Previous single-virion membrane fusion experiments have led to a comprehensive model for the mechanism by which broadly neutralizing antibodies targeting the base of HA (base bnAbs) inhibit viral membrane fusion (9, 10, 14–17). Base bnAbs bind HA in the vicinity of the fusion peptide and prevent the low pH-induced HA conformational changes (14–16). Base bnAbs thus reduce the number of activatable HAs interfacing the target membrane and either delay or prevent fusion-cluster formation (9, 10, 17). Slower fusion is inconsequential for infectivity, but lower fusion efficiency reduces the probability of productive infection (17). A greater number of unbound HAs on filamentous virions or a greater probability of membrane insertion by the unbound HAs can enable fusion in the presence of base bnAbs (17, 18). Such viral strategy that circumvents inhibitor effects and does not depend on mutations in the drug binding site can be powerful in enabling resistance to inhibitors targeting the conserved, functionally constrained epitopes.

Here we investigated the mechanism of IAV resistance to Arbidol, a broad-spectrum antiviral effective against a range of enveloped and nonenveloped viruses (12, 13, 19–23). In the context of IAV, it binds to and stabilizes the pre-fusion HA inhibiting its conformational changes at low pH (13). More specifically, Arbidol binds to a conserved cavity about 15 angstroms away from the fusion peptide where it staples the N-terminal portion of HA2 from one monomer to the central coiled-coil of an adjacent monomer(13) (Figure 1). Arbidol binding lowers the pH-threshold for HA conformational changes, likely by disfavoring HA2 coiled-coil extension. Arbidol resistance mutations, however, do not overlap its binding site, and most were shown to raise the pH-threshold of HA conformational changes (12, 13, 19). Arbidol resistance mutations overlap those of amantadine, a weak base that raises endosomal pH (11). This pattern of resistance suggests an indirect mechanism of Arbidol evasion despite its direct binding to HA, but the mechanistic picture is lacking.

**Figure 1.**
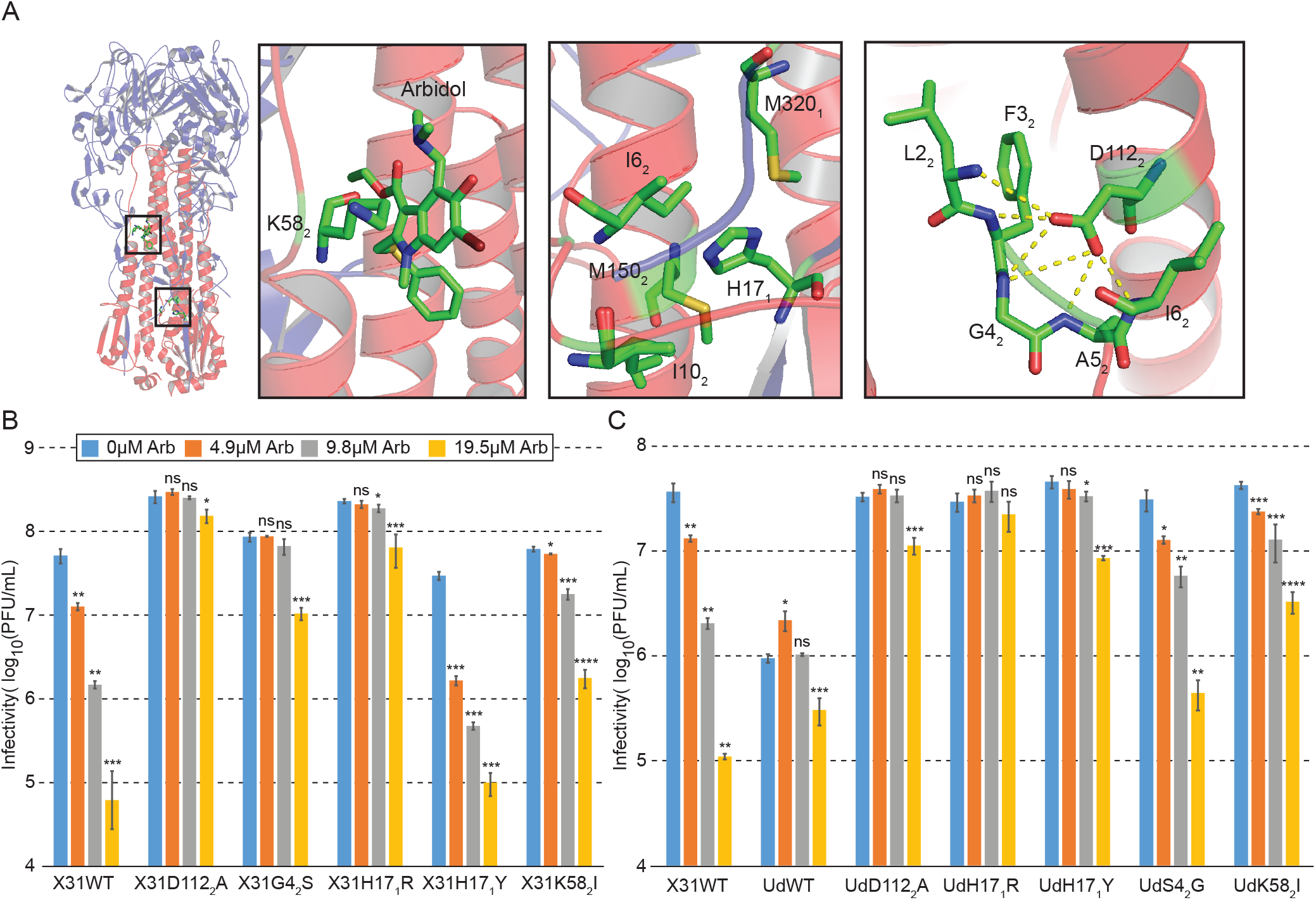
Destabilizing and stabilizing mutations confer Arbidol resistance and increased sensitivity, respectively. a) Arbidol binding site and HA region with engineered mutations are highlighted on the ribbon diagram of X31 HA in complex with Arbidol (black squares, left) (PDB: 5T6N). Bound Arbidol and each of the mutated residues are separately shown as close-ups (right). Dashed lines show non-covalent interactions within a 4Å-radius of connecting atoms. b,c) Results from the infectivity (plaque) assay with WT and mutant IAVs in the presence of varying concentrations of Arbidol. b) X31 HA and c) Udorn HA. Three biological replicates were used to derive the mean (main bars) and the standard deviation (error bars). Statistical analysis was performed using one-sided Student’s t-test comparing Arbidol-treated to untreated samples for each virus; Not significant (NS), P>0.05; *P<0.05; **P<0.01; ***P<0.001; ****p<0.0001.

We directly probed the mechanism of Arbidol resistance using our established single-virion membrane fusion platform and a panel of mutant viruses with known consequences on HA conformational dynamics during membrane fusion (4, 11, 24, 25). We found that the degree of viral sensitivity to Arbidol is a direct function of fusion-peptide stability on the pre-fusion HA. Arbidol cannot inhibit HA conformational changes when fusion peptides are sufficiently destabilized (by mutation or pH), and, consequently, it is more effective when fusion peptides are more stable. The dominance of the fusion-peptide stability over Arbidol’s effects is consistent with the notion that HA2 coiled-coil extension drives and does not follow fusion-peptide release. Our combined results thus establish a general mechanism of resistance to Arbidol independent of mutating its binding site and lead to a more detailed mechanistic picture of HA-mediated membrane fusion.

## Results

To probe the mechanism of HA resistance to Arbidol, we made a panel of viruses with mutations in HA previously shown to either stabilize or destabilize the prefusion HA structure (11, 24–26) (Figure 1A). We chose D112_2_A, G4_2_S, H17_1_Y, and K58_2_I mutations in the A/Aichi/68 (X31) HA background (Figure 1A). D112_2_A eliminates several stabilizing hydrogen-bond interactions between the carbonyl group of D112_2_ with the backbone atoms at the N-terminus of the fusion peptide buried deep within the pre-fusion HA cavity (fusion-peptide pocket) (4). G4_2_S mutation in the fusion peptide also disrupts hydrogen bonding between the fusion peptide and D112_2_ by disallowing the required fusion-peptide conformation otherwise allowed by G4_2_ (4). H17_1_ is located in a shallow hydrophobic pocked formed by M115_2_, M320_1_, I6_2_ and I10_2_ on the periphery of the fusion peptide near the HA surface. The presence of the charged R17_1_ would be unfavorable in this hydrophobic region and likely similar to what protonation of His17_1_ achieves at the pH of fusion. Y17_1_, on the other hand, would offer even greater stabilization by hydrophobic interactions and would not acquire a charge at the pH of fusion. We further included in our panel K58_2_I mutation, also reported to stabilize HA, located about 15 angstroms away from the fusion peptide adjacent to the Arbidol binding site on HA (27). We additionally generated the same panel of mutations in the context of A/Udorn/72 HA. In both cases the remaining segments derived from the A/Udorn/72 strain. Since WT Udorn HA already contains the destabilizing S4_2_, we included in its context the stabilizing S4_2_G mutation. Some of the Udorn mutants could thus be considered double destabilizing (G4_2_S/H17_1_R) or a combination of destabilizing and stabilizing mutations (e.g. G4_2_S/H17_1_Y) for probing the dominance hierarchy of mutational effects as it relates to Arbidol sensitivity (see Discussion).

We measured infectivity of our panel of mutant viruses in a standard plaque assay in the presence of a range of Arbidol concentrations from 0 to ~19.5 μM (Figure 1B-C). Arbidol treatment reduced infectivity of WT X31HA virus about 4- to 1000-fold in a concentration-dependent manner. All destabilized mutants in the X31HA background (D112_2_A, G4_2_S, and H17_1_R) were nearly completely resistant to Arbidol. A small but significant effect of Arbidol treatment was noted for only the highest Arbidol concentration, which at least in part resulted from its cytotoxicity. In fact, we could not increase Arbidol concentration in the cell-based experiments any further without causing significant cell death. Notably, the stabilizing H17_1_Y mutation resulted in an even greater sensitivity to Arbidol with close to 20-fold inhibition at 4.9 μM Arbidol, the lowest concentration tested (compare to ~4-fold inhibition of WT at this concentration). The effect of the stabilizing K58_2_I mutation did not follow the resistance pattern of the remainder of the X31HA-based panel. Significant inhibition was observed for the K58_2_I mutant at an intermediate Arbidol concentration and which had no effect on the destabilized mutants, but the mutant was inhibited less than WT. The seeming discrepancy in the resistance phenotype for the K58_2_I mutant might result from its location on HA structure adjacent to the Arbidol binding where it is likely to reduce Arbidol binding affinity directly. Udorn HA-based panel showed a similar pattern of resistance with one notable exception (Figure 1C). In accord with X31 HA results, WT Udorn was fully resistant, and this phenotype was reversed by the stabilizing S4_2_G mutation. We observed no additional effect of destabilizing mutations (D112_2_A and H17_1_R) in this context. Interestingly, the stabilizing K58_2_I mutation resulted in greater Udorn HA-virus sensitivity to Arbidol but not to the same extent as the S4_2_G mutation and resembled the phenotype of the X31HA-K58_2_I mutant. Most notably, H17_1_Y mutation in Udorn HA did not reverse the resistance phenotype of WT Udorn. We interpret H17_1_Y Udorn HA phenotype to result from the dominant effect of the destabilizing S4_2_ over the stabilizing Y17_1_, and expand upon this interpretation later (see Discussion). In sum, our results show that Arbidol sensitivity for the most part correlates with HA stability, with destabilizing mutations conferring resistance and stabilizing mutations greater sensitivity to Arbidol during infection.

We sought to distinguish between three plausible mechanisms for the observed resistance or sensitivity to Arbidol by HA mutants. One possibility was that HA destabilization compensated for Arbidol effects kinetically by increasing the overall rate of fusion even when a fraction of HAs are slowed down or unable to participate due to Arbidol binding. However, this interpretation did not seem likely in the context of our previous conclusion that rate changes during endosomal fusion are inconsequential for infectivity (17). A second possibility was that destabilizing HA mutations altered the Arbidol binding site allosterically. Finally, the destabilized HAs might permit Arbidol binding but resist its stabilizing effects at the pH of fusion.

To dissect the mechanism of Arbidol resistance for HA mutants, we performed total internal reflection fluorescent (TIRF)-based experiments of membrane fusion which permit measurements of both the rate and efficiency of membrane fusion at the single-virion level (Figure 2A). We chose two resistant (D112_2_A and H17_1_R) and the more sensitive (H17_1_Y) X31HA viruses for the detailed analyses. Viruses were preincubated with Arbidol and then allowed to bind to a supported lipid bilayer incorporating sialic-acid receptors and a pH-sensitive dye fluorescein. Fusion was triggered by introducing low-pH buffer containing Arbidol into the flow cell and monitored by fluorescence dequenching of the lipophilic DiD dye incorporated in the viral membrane. We extracted time delay for individual virions from the time of fluorescein dissipation to hemifusion (Figure 2 and Supplementary Figure 1–2). We determined hemifusion yield as the fraction of detected virions that underwent hemifusion in each field of view within fifteen minutes of observation (Figure 2D). The fifteen-minute mark was chosen as the cutoff because a clear decay in the frequency of hemifusion events was observed by this time for even the slowest of mutants at the highest Arbidol concentration suggesting that measurements approached true hemifusion efficiency in all cases (Supplementary Figure 1 and 2). In the presence of Arbidol, hemifusion lag-time distributions at pH5.2 shifted toward slower times for the WT, but remained unchanged for the destabilized mutants (Figure 2B-D and Supplementary Figure 1). Furthermore, in the presence of Arbidol, hemifusion yield was lower for WT, but remained unchanged for the destabilized mutants (Figure 2D). The decrease in hemifusion efficiency for WT virus at pH5.2 is consistent with Arbidol preventing rather than slowing down the conformational change of bound WT HAs (4, 17). The unchanged rate of fusion for the destabilized mutants confirms true resistance rather than kinetic compensation by the destabilized HAs (Figure 2C). Together, our single-virion fusion experiments showed that Arbidol prevents the low pH-induced conformational changes of WT HA, and HA destabilization confers resistance to Arbidol.

**Figure 2.**
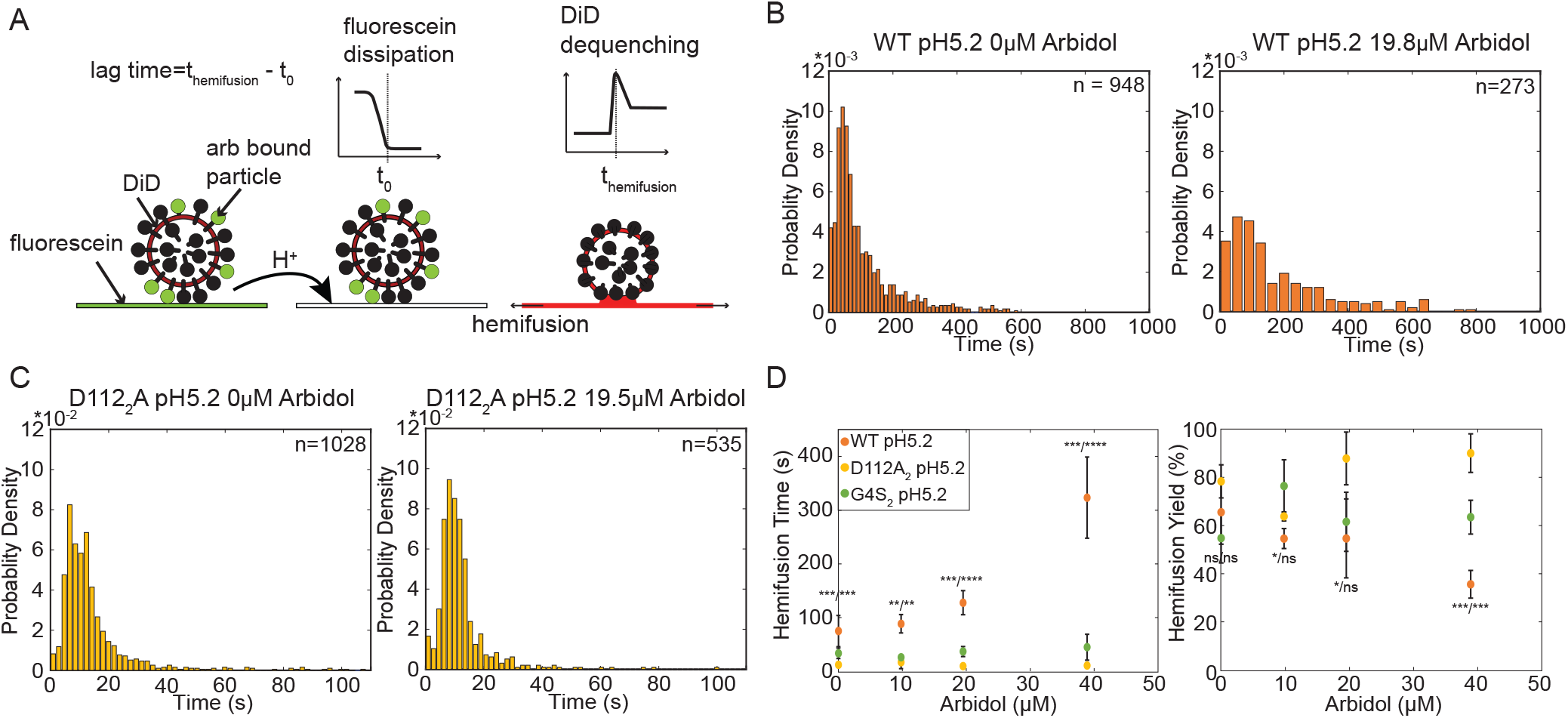
Destabilizing HA mutations confer Arbidol resistance to IAV in a TIRF-based single-virion membrane-fusion assay. a) Schematic of the experiment setup. Virions are pre-incubated with Arbidol and bound to the supported lipid bilayer via sialic-acid receptors displayed on GD1a. Fluorescein on the target membrane serves as the pH-sensor. DiD is incorporated into the viral membrane so that it is partially quenched. DiD dequenching indicates hemifusion between the viral and the target membranes. Hemifusion time is measured as the delay from fluorescein dissipation to DiD dequenching for individual virions. Hundreds to thousands of virions are recorded in each field of view. b,c) Hemifusion lag time probability-density histograms for WT (b) and D1122A (c) in the presence or absence of 19.5 μM Arbidol. Single-virion data are pooled from different experiments and fitted with Gamma distribution (black line). The total number (n) of virions represented by histograms is indicated on each plot. d) Median hemifusion lag time (left) and yield (right) for WT, G42S, and D1122A derived from individual experiments are plotted as the average ± standard deviation. Statistical analysis was performed using one-sided Student’s t-test at each Arbidol concentration. The P values comparing each D1122A and G42S to WT are indicated with symbols as P(D1122A) / P(G42S). Not significant (NS), P>0.05; *P<0.05; **P<0.01; ***P<0.001; ****p<0.0001.

Our previous single-virion membrane fusion experiments related the pH-dependence of the fusion rate to the probability that HA2 extends while HA1 remains in the open state (4). In this model, the probability of HA2 extension is related to the stability of the fusion peptide in its pre-fusion pocket, and the time that HA1 spends in the open state is determined by pH. The pattern of sensitivity and resistance to Arbidol by WT and mutant viruses (Figure 2 and Supplementary Figure 1) suggests that Arbidol delays WT but not destabilized HA extension past the window of opportunity provided by HA1 opening. The destabilized HAs might nonetheless be bound by Arbidol, but Arbidol might not prevent their extension at low pH. To probe whether HA destabilization is sufficient for allowing evasion of Arbidol’s effects, we performed single-virion membrane fusion experiments with WT X31 HA virus in the presence of Arbidol at pH 4.8. We chose pH 4.8 because it represents the threshold pH, below which the rate of WT X31HA conformational changes is no longer pH-dependent and thus no longer limited by HA1 opening (4, 8). Furthermore, the hemifusion rate of WT IAV below pH 4.8 approximates that of destabilized mutants, suggesting that pH in this range destabilizes fusion peptides to a similar extent as the destabilizing mutations, and/or it destabilizes them sufficiently so that a different HA transition becomes rate limiting (4). We found that WT X31 HA virus resists inhibition by Arbidol at pH 4.8 and displays the pattern of resistance indistinguishable from that of destabilized mutants at pH 5.2 (Figure 3A, and compare to Figure 2D). Since WT HA is bound by Arbidol at neutral pH, this result is consistent with our interpretation that HA destabilization is sufficient to explain resistance to Arbidol (Figure 3A). To verify that the observed resistance is due to HA destabilization directly and not due to reduced Arbidol affinity for the pre-fusion HA at the lower pH, we performed single-virion fusion experiments with the stabilized H17_1_Y X31HA mutant virus at pH 4.8. Hemifusion by H17_1_Y X31HA mutant virus at pH 4.8 approximates that of WT X31 HA virus at pH 5.2 in both rate and efficiency (17) and displays a similar degree of sensitivity to Arbidol as WT virus at pH 5.2 (compare Figures 3A and 3B). Arbidol’s affinity for the pre-fusion HA is thus unaffected by the lower pH, and resistance of WT virus to Arbidol at pH 4.8 is owed to HA destabilization directly. Our combined results offer direct support for and extend the fusion model we proposed previously (4) (see Discussion). Furthermore, our results reveal a mechanism of drug resistance deriving from a functional modification of its target rather than from mutation of the drug-binding site.

**Figure 3.**
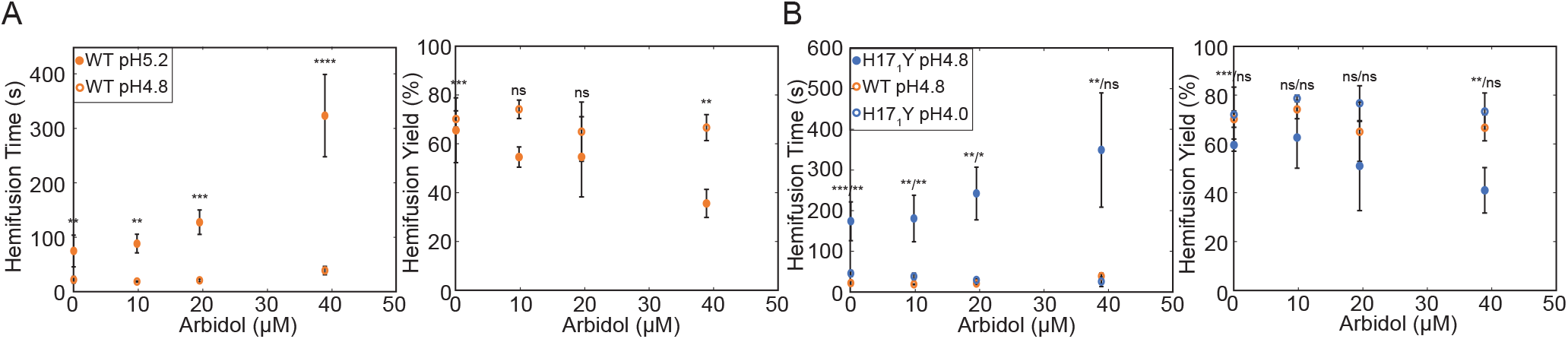
Sensitive viruses resist inhibition by Arbidol at sufficiently low pH. a) Median hemifusion lag time (left) and yield (right) for the WT X31HA virus at pH 5.2 or 4.8 are shown as the average ± standard deviation. Statistical analysis was performed using one-sided Student’s t-test comparing pH5.2 to pH 4.8 data at each Arbidol concentration. b) Median hemifusion lag time (left) and yield (right) for WT X31HA virus at pH 4.8, and the H171Y mutant virus at pH4.8 and pH 4.0, are shown as the average ± standard deviation. Statistical analysis was performed using one-sided Student’s t-test, comparing pH 4.0 to pH 4.8 data for H171Y mutant at each Arbidol concentration.

## Discussion

The current gold standard in antiviral inhibitor design is to target the conserved, functionally constrained epitopes on essential viral targets. However, mechanisms that modulate viral functions in ways that render inhibitor effects inconsequential for infectivity would represent a powerful evolutionary strategy ultimately forcing us to rework our own design standards. Our experiments with IAV and Arbidol have identified one such viral evasion strategy. Arbidol binds to a conserved site on HA and prevents HA conformational changes. Destabilization of HA fusion peptides by mutation or pH renders viruses resistant to Arbidol’s effects. The resistance phenotype is a direct function of fusion-peptide stability, and full resistance is acquired even when binding of Arbidol to the pre-fusion HA is unaltered (Figure 3). The resistance mechanism we uncovered is consistent with the observations that the resistance mutations (both published (12, 19) and the ones newly studied here in Figure 1) are not localized to a specific site on HA. It further explains the large degree of overlap between Arbidol and amantadine resistance mutations (11, 19). In the case of amantadine, HA destabilization permits membrane fusion at the raised endosomal pH. In the case of Arbidol, HA destabilization offsets stabilization by Arbidol. This model further predicts that Arbidol and amantadine resistance mutations would not have a complete overlap, but only in those HA regions that are dominant to the stabilization afforded by Arbidol binding. Future HA mutagenesis will explore this question further.

The question of HA conformational sequence at low pH has been a matter of debate in the field. Classic work by Carr and Kim led to the “spring-loaded” model of HA-mediated membrane fusion whereby the release of HA1 clamp on HA2 in open HA allows HA2 extension driven by coiled-coil extension (Loop B-to-helix transition in Figure 4A) (28). This notion was supported by experiments showing that inhibition of HA1 opening by engineered disulfide bridges prevents fusion-peptide exposure and inhibits membrane fusion (3). However, recent biophysical and theoretical explorations have brought that notion into question and suggested that fusion-peptide release might precede HA1 opening and coiled-coil extension (29–34). Our current experiments offer insight into this question by coupling interpretations of HA conformational dynamics to membrane fusion and can thus reveal a functionally relevant pathway. We found that destabilization of the fusion peptide can offset HA stabilization by Arbidol. Arbidol links Loop B from one HA2 monomer to C helix of an adjacent monomer where it must interfere with coiled-coil extension (13) (Figure 1A and illustrated in Figure 4B). If the release of the fusion peptide preceded and did not depend on coiled-coil extension, destabilization of the fusion peptide would have no bearing on HA stabilization by Arbidol. In other words, if Arbidol prevented HA2 extension on WT virus even after fusion-peptide release, faster release of the fusion peptide for the mutants or at the lower pH would not change that property. The simplest interpretation of our membrane-fusion experiments is captured by the following model (Figure 4). The probability of HA2 extension is related to the time window afforded by HA1-head opening and the composite of two critical forces: the force exerted on the fusion peptide by the coiled-coil extension, and the stabilizing interactions in the fusion-peptide pocket resisting fusion-peptide release. Arbidol binding incurs free-energy penalty for coiled-coil extension and prevents extension of WT HA at the pH of fusion. Fusion-peptide destabilization by mutation or pH reduces free-energy penalty for fusion-peptide release and permits HA2 extension in the presence of Arbidol. Our conclusions thus support the “spring-loaded” model of HA-mediated membrane fusion and contrast those depicting fusion-peptide release before HA1 opening and coiled-coil extension (29-32, 34). Reconciling these contrasting results, it is possible that fusion-peptide release before HA1-head opening might lead to nonproductive refolding and HA inactivation. Indeed, our previous analysis of HA mediated membrane fusion revealed that about half the HAs interfacing the target membrane become inactivated instead of inserting into the target membrane over the course of the fusion reaction (10). To distinguish productive and nonproductive HA-refolding pathways, analyses of HA conformational dynamics must be coupled to measurements of membrane insertion or fusion.

**Figure 4.**
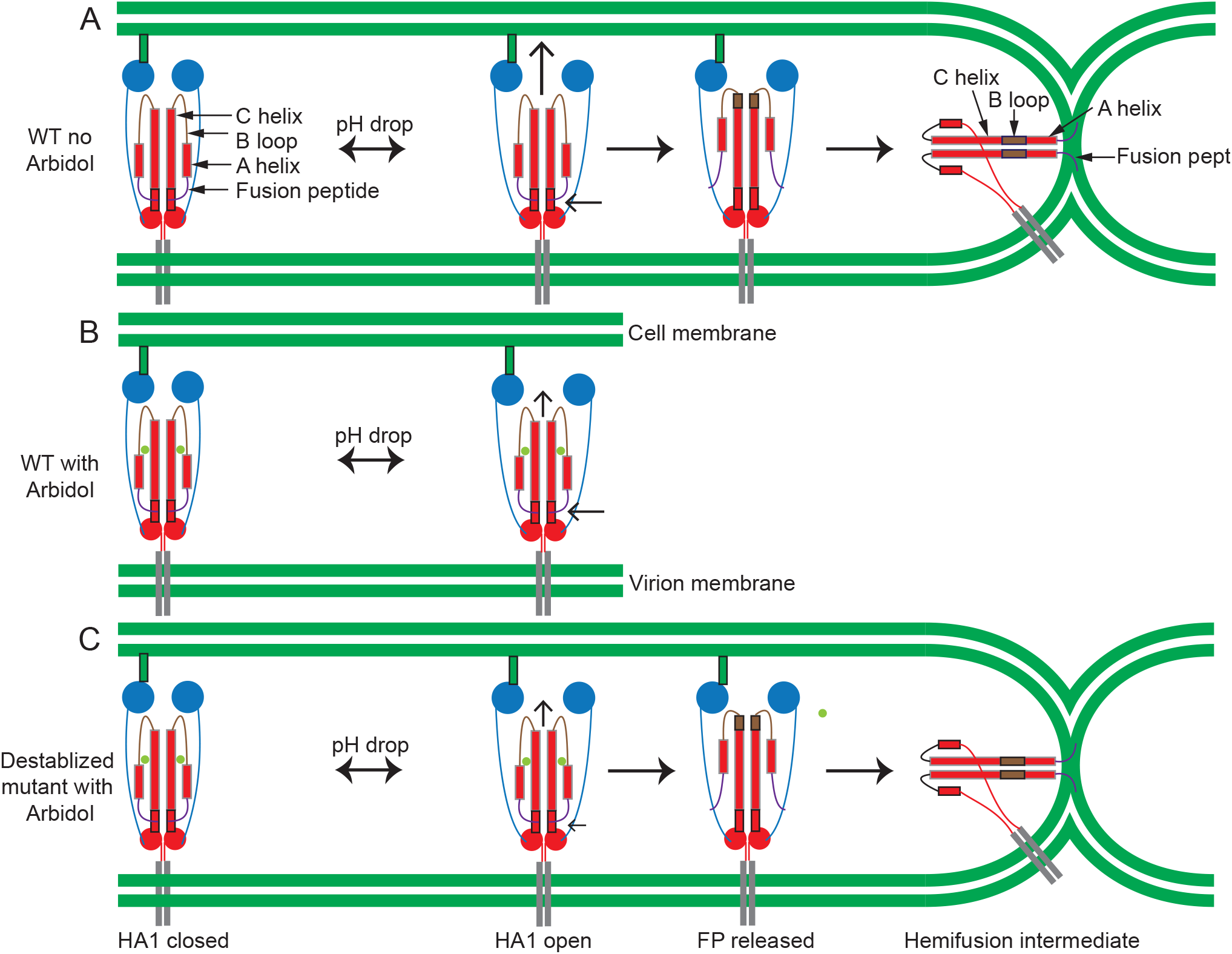
An updated model of HA-mediated membrane fusion and the mechanism of Arbidol resistance. a) Low pH increases the average time that HA spends in the open state. The probability of HA2 extension is related to the free energy released by coiled-coil extension (loop B-to-helix transition; the tug, upward arrow) and the stability of the fusion peptides in their pockets (resistance, side arrow) during the window of opportunity afforded by HA1-head separation. B) Arbidol binding to HA links loop-B residues on one HA2 monomer to C-helix residues on an adjacent monomer. It prevents HA2 extension by incurring free-energy penalty to coiled-coil extension (smaller upward arrow). c) Fusion-peptide destabilization by mutation or pH lowers the free-energy penalty for its release (smaller side arrow) allowing coiled-coil extension despite Arbidol binding to pre-fusion HA.

Infectivity experiments with A/Udorn/72 mutants allowed us to probe the dominance hierarchy of mutational effects as it relates to Arbidol sensitivity. WT Udorn is resistant to Arbidol owing to its destabilizing S4_2_ (Figure 1C). Indeed, Udorn S4_2_G is sensitive to Arbidol and resembles X31 WT in both sensitivity (Figure 1C) and membrane fusion kinetics (4). H17_1_Y mutation has different consequences in the two WT backgrounds. In the context of G4_2_ on A/Aichi/68, H17_1_Y is strongly stabilizing ((17, 25) and Figure 3B) and confers greater sensitivity to Arbidol (Figure 1B, 3B, and Supplemental Figure 2). However, H17_1_Y in the context of S4_2_ on A/Udorn/72 did not diminish Arbidol resistance (Figure 1C) suggesting that the destabilizing effect of S4_2_ is dominant over the stabilizing effect of Y17_1_. The main region conferring fusion peptide stability thus resides deep in its binding pocket.

Base bnAb also bind the pre-fusion HA and inhibit conformational changes at the pH of fusion (14–16). However, destabilization of the fusion peptides does not confer resistance in this case (17). Greater effectiveness of base bnAbs might be owed to their larger footprint or their binding in the immediate periphery of the fusion peptides or both (14–16). Our work has thus identified a potential limitation of targeting the pre-fusion HA by small molecules. More work is needed to determine whether mutations of the type identified here could become relevant in clinic. Fusion peptide and its pocket on pre-fusion HA represent the most conserved, signature feature of HA across subtypes. It remains an open question whether this conservation represents a functional requirement in the wild or simply reflects the current absence of pressure on this highly concealed HA region.

## Materials and Methods

### Reagents

#### Cells

MDCK.2 (ATCC strain CCL-34) and Human Embryonic Kidney 293T (HEK293T) cells (ATCC strain CRL-3216) were propagated in DMEM supplemented with 10% FBS in humidified 37°C incubator with 5% CO_2_. All infections were performed in infection media (OptiMEM (Thermo Fisher Scientific) supplemented with 1μg/ml TPCK-trypsin (Sigma-Aldrich)). 6-Bromo-4-((dimethylamino)methyl)-5-hydroxy-1-methyl-2-((phenylthio)methyl)-1H-Indole-3-carboxylic acid ethyl ester monohydrochloride (Arbidol) (Sigma Aldrich) was dissolved in ethanol at 10mg/ml and diluted to target concentration in infection media.

#### Viruses

IAV used in this study have HA from either A/Aichi/68(X31) or A/Udorn/72(Ud) and the remaining segments from Ud (35). Viruses were propagated and purified as described previously (17). In brief, viruses were passaged at a multiplicity of infection (MOI) of 0.001 PFU/cell except for H17_1_Y, which was passaged at an MOI of 0.1. Viruses were passaged twice at the low MOI before a high-MOI infection of 12 PFU/cell and virus purification. After each infection step, the infected-cell supernatants were clarified by centrifugation at 1000xg for 10min. After the final, high-MOI infection, the clarified supernatants were centrifuged through a 20% sucrose cushion in HNE20 (20mM HEPES, 150mM NaCl, 0.2mM EDTA, pH 7.4) at 100,000xg for 2.5 hours at 4°C to concentrate and partially purify the virus. The viruses were then further purified by centrifugation through a 20-60% (w/v) sucrose gradient at 100,000g for 2.5h. The prominent top band, enriched in spherical virions was collected (4, 17). Second-passage viruses were used in plaque reduction assays (Figure 1) and purified spherical viruses were used in single-virion hemifusion experiments (Figures 2–3 and Supplementary Figures 1–2).

### Infectivity experiments

Second-passage viruses of indicated strains were preincubated with infection media with 0, 4.5, 9.8, or 19.5 μM Arbidol at room temperature for 1 hour before virus attachment to confluent MDCK.2 cell monolayers. Virus concentration in pre-incubations was adjusted to about 2×10^10^-2×10^11^ PFU/ml. Virus was then diluted ten-fold serially in infection media containing the same Arbidol concentration, and 100ul of virus dilutions was added per well of a 6-well plate. Attachment was performed at room temperature for 1 hour with shaking every 6 min. After attachment, cells were overlayed with OptiMEM containing 1ug/ml TPCK-Trypsin, 0.6% Oxoid Agar (Themo Fisher Scientific), and the same Arbidol concentration used in pre-incubations. The plates were incubated at 34°C for two days before fixing with 3.7% formaldehyde in phosphate buffered saline (PBS) (137mM NaCl, 2.7mM KCl, 10mM Na_2_HPO_4_, 1.8mM KH_2_PO_4_ pH7.4) at room temperature for 30min. Overlayed media was removed, monolayers washed with PBS, and the plate was imaged in brightfield using Keyence fluorescent microscope BZ-X800 (Keyence) equipped with a 2x objective (Keyence Plan Apochromat, 0.1 NA). Light intensity was adjusted to 25% and exposure to 1/3,000 seconds. Wells containing 20-200 plaques were scanned, and separate images corresponding to different parts of the well were stitched together using BZX-800 Analyzer software (version 1.1.2.4) to assemble a full-well image version for counting. The experiment including an entire panel of viruses was repeated at least three times. Figures 1B and 1C show a representative result of an experiment performed in three biological replicates.

### Single-virion membrane fusion experiments

#### Hemifusion Assay

Experiment followed an established procedure with only slight modifications to include Arbidol (pre)treatment of virions (17). In brief, 6μg of purified virus was labeled with 1,1′-dioctadecyl-3,3,3′,3′-tetramethylindodicarbocyanine, 4-chlorobenzenesulfonate salt (DiD, Thermo Fisher Scientific) at 10 μM for 1.5 h at room temperature in a 25μl reaction. Liposomes consisted of 4:4:2:0.1:2 × 10−4 ratio of 1,2,dioleoyl-sn-glycero-3-phosphocholine (Avanti Polar Lipids), 1-oleoyl-2-palmitoyl-sn-glycero-3-phosphocholine (Avanti Polar Lipids), sn-(1-oleoyl-2-hydroxy)-glycerol-3-phospho-sn-3′-(1′-oleoyl-2’-hydroxy)-glycerol (ammonium salt) (18:1 BMP (R,R); Avanti Polar Lipids), bovine brain disialoganglioside GD1a (Sigma-Aldrich) and N-((6-(biotinoyl)amino)hexanoyl)-1,2-dihexadecanoyl-sn-glycero-3-phosphoethanolamine (biotin-X DHPE; Molecular Probes, Life Technologies). Planar bilayers were formed from 200-nm liposomes in channels of a PDMS flow-cell using vesicle spreading method (36). Sialic acid on GD1a served to attach IAV virions to the bilayer. Fluorescein-conjugated streptavidin (Invitrogen) at 30 μg/ml was bound to the bilayer and served as a pH-indicator. DiD-labeled virions (untreated or pre-treated with Arbidol at room temperature for 30min) were flowed into the channels and allowed to attach at room temperature for 10min. Unbound virions were washed out using HNE20, and pH was dropped by flowing in the low-pH buffer at the indicated pH (10mM Citrate, 140mM NaCl, 0.1mM EDTA). The wash and the low-pH buffers contained Arbidol concentration matching the pre-incubation conditions. The flow of the low pH buffer was adjusted to 60μl/min for the first minute, and then to 20μl/min for the rest of the experiment. Hemifusion was monitored as DiD dequenching from individual virions. All experiments were perfomed at 23±0.5°C ambient temperature.

#### Microscope Configuration

The excitation and imaging pathways were unchanged relative to our previous set-up except for the camera (17). Some of the movies were imaged using the original 512 × 512 pixel EM-CCD sensor (Model C9100-13; Hamamatsu), and some using a newer, 1024 x 1024 pixel EM-CCD sensor (Model C9100-24B; Hamamatsu). 488-nm laser at 1-2μW (Obis, Coherent) was used to excite fluorescein, and 647-nm laser at 0.5-1μW (Obis, Coherent) was used to excite DiD. The exposure time was 0.4 or 0.5 seconds depending on virus and Arbidol concentration.

### Data analysis and statistics

All data analysis for single-virion experiments was performed using MATLAB (MathWorks). pH-drop time and DiD-dequenching time for individual virions were derived as described previously (4, 17). We determined hemifusion yield as the percentage of detected virions that hemifused within 15 minutes of the pH-drop. Statistical analyses for infectivity measurements, hemifusion time and hemifusion yield were performed using one-tailed unpaired t-tests, assuming equal variances. The *P* values are designated using the following symbols: NS, *P* > 0.05; \**P* < 0.05, the null hypothesis was rejected at the 5% significance level; \*\**P* < 0.01, the null hypothesis was rejected at the 1% significance level; \*\*\**P* < 0.001, the null hypothesis was rejected at the 0.1% significance level; \*\*\*\**P* < 0.0001, the null hypothesis was rejected at the 0.01% significance level.

## Acknowledgments

We thank Stephen C. Harrison (Harvard Medical School) for comments on the manuscript. We acknowledge support from the NIH Director’s New Innovator Award 1DP2GM128204 (to T.I.) and the NSF MRSEC DMR-1420382 (to T.I.).

---

**Supplemental Figure 1.**
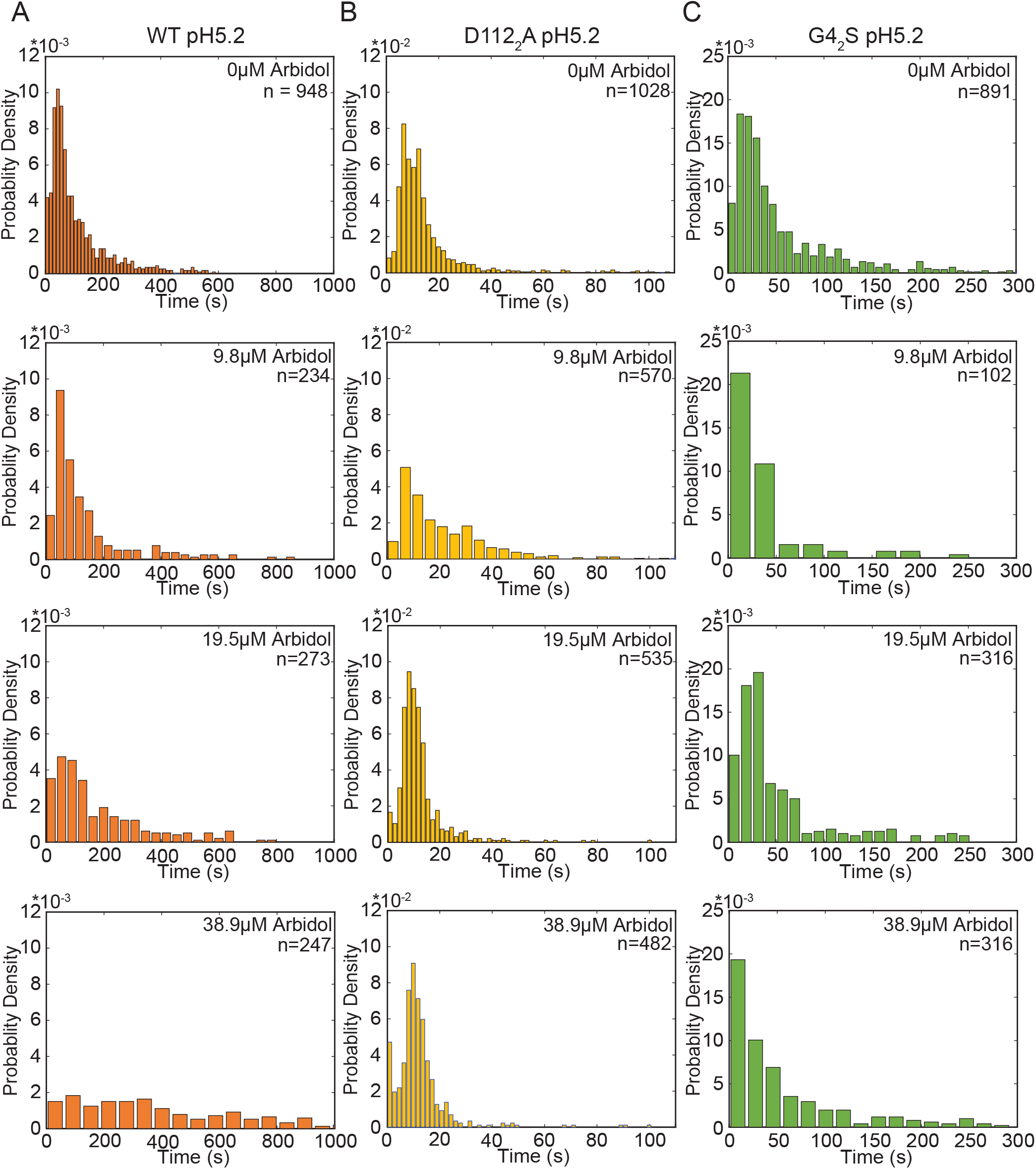
Hemifusion lag time probability-density histograms for WT (a), D1122A (b) and G42S (c) X31HA IAV at pH 5.2 in the presence of indicated Arbidol concentrations.

**Supplemental Figure 2.**
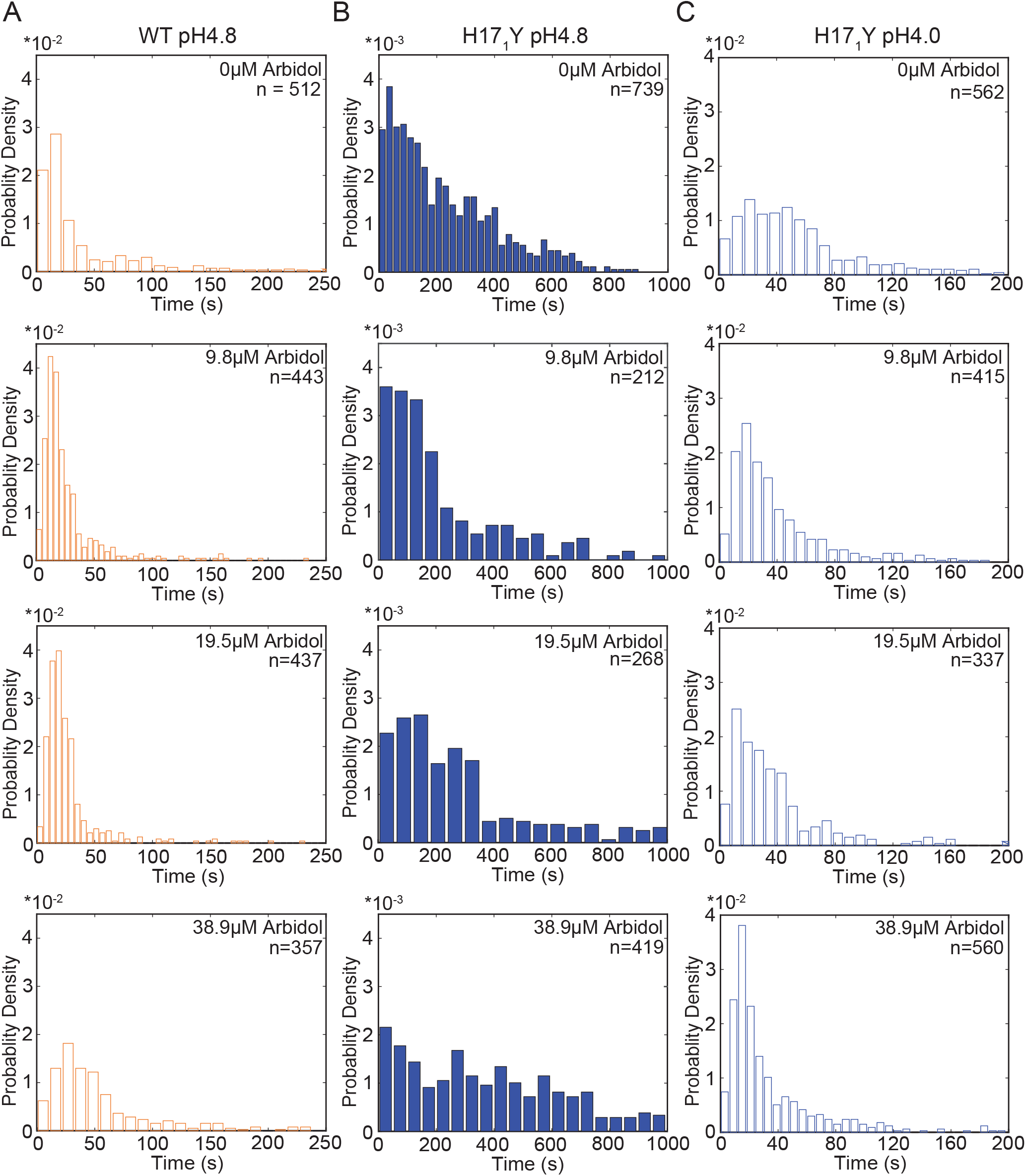
Hemifusion lag time probability-density histograms for H171Y pH 4.8 (a) WT pH 4.8 (b) and H171Y pH 4.0 (c) X31HA IAV in the presence of indicated Arbidol concentrations.

